# Disentangling motor and cognitive roles of cortical beta oscillations and their modulation with rTMS

**DOI:** 10.64898/2025.12.08.692972

**Authors:** Shenghong He, Juan Francisco Martin-Rodriguez, Alek Pogosyan, Eduardo M. Moraud, Huiling Tan

## Abstract

Cortical beta oscillations are central to motor control. These rhythms are modulated during movement preparation and execution, and are also sensitive to cognitive factors such as uncertainty, attention and reward prediction. However, the distinct motor and cognitive roles of cortical beta oscillations —roles that may inform condition-specific therapeutic strategies—remain unclear. This study aimed to disentangle the respective contributions of motor and cognitive functions, and to assess their modulation with repetitive transcranial magnetic stimulation (rTMS). Twenty-four healthy participants performed a visually cued reaching task in which directional uncertainty was dynamically manipulated by varying the number of targets in real-time. Reaction time was quantified as a behavioural measure. Three stimulation conditions were applied during the preparatory periods between the ‘uncertainty’ and ‘go’ cues: no stimulation, regular rTMS at each participant’s individual beta frequency, and irregular rTMS. Electroencephalography was used to measure beta-band oscillatory activity. Our results showed that ‘uncertainty’ cues induced bilateral beta suppression, with greater uncertainty linked to smaller reductions in beta power. Movement-related beta modulation was lateralised to the hemisphere contralateral to the executing hand, where elevated beta power in the pre-‘go’ cue period predicted longer reaction times. Both regular and irregular rTMS significantly shortened reaction times and attenuated beta event-related desynchronisation. Reductions in beta desynchronisation correlated with improvements in reaction times, suggesting beta desynchronisation reflects a neural threshold for movement initiation. These findings indicate that cortical beta oscillations encode distinct motor and cognitive processes, and that beta frequency rTMS can modulate beta dynamics to facilitate faster movement initiation by lowering this neural threshold.

## Introduction

Beta-band oscillations constitute a prominent feature of the cortical-basal ganglia motor network, and are dynamically modulated by movements across cortical and subcortical structures (**McFarland et al., 2000; Pfurtscheller et al., 2003**). Typically, movement preparation is accompanied by a gradual beta event-related desynchronisation (ERD), with beta power decreasing and reaching a minimum around movement initiation. This is followed by an event-related synchronisation (ERS) after movement termination, commonly referred to as the ‘beta rebound’ (**Pfurtscheller and Lopes da Silva, 1999; Zaepffel et al., 2013**). Beyond their role in movement preparation and execution, cortical-basal ganglia beta rhythms are also influenced by cognitive aspects of motor control. For instance, beta amplitude has been shown to increase during anticipation of an informative cue, peaking around cue onset (**Saleh et al., 2010**). Moreover, beta suppression during movement preparation is modulated by uncertainty about the direction of the upcoming movement (**Tzagarakis et al. 2010**). Collectively, these findings support the view proposed by Jenkinson and Brown that beta-band activity arises from multiple sources and reflects distinct functional processes—a concept they termed “functional polymorphism” (**Jenkinson and Brown, 2011**). In addition, Engel and Fries suggested that beta activity reflects the maintenance of the current sensorimotor or cognitive state, serving as a neural correlate of the “status quo” (**Engel and Fries, 2010**). Despite these consensuses, the distinct characteristics of the movement-related versus cognition-related beta oscillations have yet to be clearly delineated. A clearer dissociation between these two functions of beta activity is crucial to elucidate how neurodegenerative disorders perturb the balance between movement execution and higher-order control, and to inform the next generation of neuromodulation strategies.

In people with Parkinson’s disease (PD), exaggerated beta-band activity in the cortical-basal ganglia motor circuit is a hallmark of the disease pathophysiology and contributes to motor impairments (**Brown, 2007**). This dysregulation is accompanied by alterations in the beta ERD/ERS pattern as well (**Leocani & Comi, 2006**), with the onset latency of beta ERD shown to correlate with mean reaction time (**Kühn et al., 2004**). Therapeutic interventions such as L-dopa administration or deep brain stimulation (DBS) of the subthalamic nucleus (STN) have been found to restore both beta oscillatory dynamics and motor performance (**Devos & Defebvre, 2006; Mathiopoulou et al., 2024**). Notably, improvements in motor symptoms following medication or DBS are positively associated with suppression of STN beta power (**Kühn et al., 2006; Neumann et al., 2016; Kehnemouyi et al., 2021**). These findings have motivated the development of beta-triggered adaptive DBS (aDBS) strategies (**Little et al., 2013**), with a recent international clinical trial (NCT04547712) demonstrating that long-term aDBS is tolerable, effective, and safe in PD patients previously stabilised on continuous DBS (cDBS) (**Bronte-Stewart et al., 2025**).

In parallel, various non-invasive neuromodulation techniques—such as neurofeedback training, repetitive transcranial magnetic stimulation (rTMS), and transcranial direct/alternating current stimulation (tDCS/tACS)—have been investigated for their potential to modulate cortical beta oscillations in both healthy individuals and people with PD (**Chou et al., 2015; Duan and Zhang, 2024; He et al., 2020a, 2020b; Pogosyan et al., 2009**). While these approaches have shown promise in influencing neural dynamics and motor behaviour, their electrophysiological and behavioural impact remain inconsistent, and the mechanisms through which they act are not yet fully understood. This variability has so far limited their translational value.

To address these limitations, here we aimed to disentangle the motor and cognitive roles of cortical beta oscillations within a unified experimental framework, including movement preparation and execution under varying levels of directional uncertainty. Additionally, we investigated how these oscillations and their associated behavioural outcomes are modulated by distinct rTMS paradigms. Participants were asked to perform a reaching task, in which trial-by-trial reaction time was quantified as the behavioural readout. Directional uncertainty was dynamically manipulated during movement preparation by varying the number of potential targets before the ‘go’ cue. Three stimulation conditions were applied during the preparatory period between the ‘uncertainty’ cue, and the ‘go’ cue: no stimulation, regular rTMS at the individual’s beta frequency (IFB), and irregular rTMS.

## Materials and Methods

### Participants

Twenty-four healthy human subjects participated in the study (twelve males and twelve females; age range 19-33 years; all right-handed). The study was approved by the local Research Ethics Committee (Med IDREC Ref: R35227/RE001) and performed in accordance with the Declaration of Helsinki, and strictly following the international safety guidelines for TMS (**Rossi et al., 2009**).

### Determination of individual beta frequency (IBF)

For each participant, the IBF was determined prior to the experiment following the procedure described by **Romei et al. (2016)**. Briefly, EEGs were recorded while the participant was performing a simple self-paced right index finger tapping task on the space bar of a computer keyboard (**Fig. 1C**), at a pace of approximately 0.25 Hz for a total of 45 repetitions. The IBF was identified from EEGs recorded over the left primary motor cortex (electrode C3) as the frequency within beta range (13-30 Hz) that exhibited the maximum post-movement beta rebound. On average, the IBF was 20.44 ± 0.38 Hz (SEM), with a range of 18-23 Hz.

**Fig. 1.**
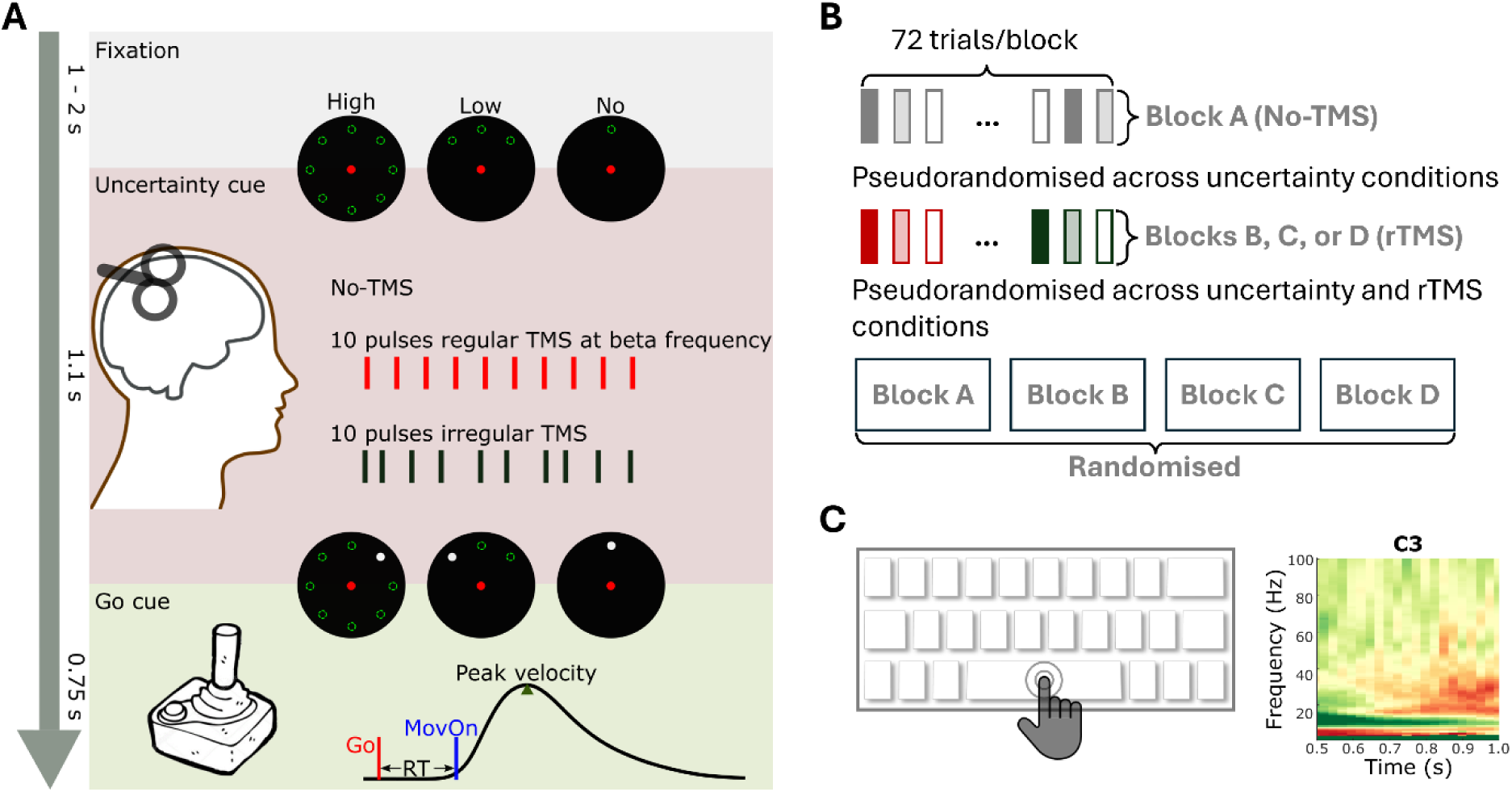
Experimental paradigm. **(A)** Timeline of a single trial involving a sequence of fixation, preparation with ‘uncertainty’ cue, rTMS delivery (three possible conditions), ‘go’ cue, and reaching task. **(B)** The experiment consists of four blocks of 72 trials each, with blocks and trials presented in a randomised or pseudorandomised order. **(C)** Prior to the experiment, each participant’s individual beta frequency is determined as the frequency within the beta frequency range (13-30 Hz) that shows maximal event-related synchronisation during a button-clicking task (**Romei et al., 2016**).

### Experimental paradigm

Each participant was invited to perform a delayed reaching task. As shown in **Fig. 1A**, each trial begins with a ‘fixation’ period lasting 1 – 2 seconds, during which the participant is instructed to focus on the screen. This is followed by a ‘preparatory phase’ with a cue of high, low, or no directional uncertainty, modulated by varying the number of potential targets displayed as green dashed hollow circles. A high ‘uncertainty’ cue presents eight potential targets evenly distributed across 360°, while a low ‘uncertainty’ cue displays three potential targets spanning 90°. In contrast, a no ‘uncertainty’ cue includes only the target located at the top-centre of the screen. The last phase of each trial is the ‘go phase’ with the ‘go’ cue, appearing as a white solid circle, presented 1.1 seconds after the ‘uncertainty’ cue. Upon its appearance, the participant is instructed to use a joystick to move from the starting position (indicated in red) to shoot the target (indicated in white) as quickly and accurately as possible. Within 0.75 seconds of the ‘go’ cue, accuracy feedback is provided via an animation, i.e., if the movement endpoint falls within 5° of the target, the target ‘exploded’, indicating a successful hit. An inter-trial interval of 2.75 second is included before the onset of the next trial. Between the ‘uncertainty’ cue and the ‘go’ cue, there are three rTMS conditions including no TMS, regular rTMS, and irregular rTMS. In the regular rTMS condition, 10 TMS pulses are delivered at the participant’s IBF. In the irregular rTMS condition, 10 TMS pulses are delivered at pseudorandom, unequally spaced intervals, with the timing of the last pulse with respect to the first pulse and (i.e., total duration between the first and last pulses) matched to that of the regular rTMS condition (e.g., 0.5 seconds for an IBF of 20 Hz). The onset of the first rTMS pulse varies randomly between 0.35-0.55 seconds after the ‘uncertainly’ cue. As shown in **Fig. 1B**, the experiment consists of four blocks of 72 trials each, including one no-TMS block and three rTMS blocks, presented in a randomised order. Within each block, trials from different conditions are pseudorandomised to ensure an equal number of trials per condition–three uncertainty conditions in the no-TMS block, and six combined uncertainty and rTMS conditions in the rTMS blocks.

### Data acquisition

The experimental paradigm was implemented in Python using the open-source software PsychoPy (v1.74). For each trial, PyschoPy generated two signals. The first signal contains two discrete amplitude levels, indicating the onset of two types of rTMS (either regular or irregular), and was input to a CED1401 (Cambridge Electronic Design, UK) to control the delivery of TMS pulses (Magstim Rapid, MRS1000/50, Magstim Company, UK). The second signal contains variable amplitudes encoding the timing of different task cues (‘Fixation’, ‘Uncertainty’, and ‘Go’) as well as the uncertainty levels. This signal was sampled using a TMSi Porti amplifier (TMS International, Netherlands), which was also used to record EEG, EOG, EMG, and joystick position data. Scalp EEGs were recorded from 23 channels—‘F7’, ‘F3’, ‘Fz’, ‘F4’, ‘F8’, ‘FC5’, ‘FC1’, ‘FC2’, ‘FC6’, ‘C3’, ‘Cz’, ‘C4’, ‘CP5’, ‘CP1’, ‘CP2’, ‘CP6’, ‘P3’, ‘Pz’, ‘P4’, ‘POz’, ‘O1’, ‘Oz’, and ‘O2’—according to the international 10–20 EEG system. EOGs were recorded from the left eye, and EMGs were recorded from three muscles in the right forearm/hand: the first dorsal interosseous muscle (FDI), flexor, and extensor muscles. Joystick positions were recorded as x and y coordinates relative to the centre position. All signals were sampled at 2048 Hz. EEGs were recorded in a unipolar configuration with a common average reference, while EOGs and EMGs were recorded in a bipolar configuration. Joystick positions were acquired via the amplifier’s auxiliary port.

### Determination of active motor threshold (AMT)

The TMS coil was positioned tangentially to the scalp and oriented at 45° angle from the sagittal midline to induce a posterolateral-anteromedial current flow. The optimal stimulation site (‘hotspot’) for eliciting movement evoked potentials (MEPs) in the right FDI muscle was identified and marked. This was achieved by delivering suprathreshold single-pulse TMS over the left primary motor cortex (M1) and locating the coil position that produced the largest MEP in the right FDI (**Rossini et al. 2015**). The active motor threshold (AMT) was then determined as the minimum stimulation intensity required to evoke a peak-to-peak MEP of at least 200 µV while the participant was maintaining a voluntary contraction of the target muscle at 20% of their maximum force. On average, the AMT was 51.81 ± 2.30% (SEM) of the maximal stimulator output (range: 34-66%). For each participant, the stimulation intensity for rTMS was set at 90% of their individual AMT. A Fastrack®–Polhemus neuronavigation system, controlled by custom scripts, was used to monitor and maintain consistent positioning over the TMS ’hot spot’ throughout the experimental session.

### Behavioural data analysis

For each individual trial, a reaction time (RT) was derived from the joystick position trajectories. Specifically, the Euclidean distance from the starting point was computed for each pair of XY coordinates using the square root of the sum of squares (SRSS). The distance trajectories from individual trials were visually inspected. Trials were excluded if they showed movement before the ‘go’ cue, contained obvious noise, or failed to initiate movement within 750 ms. For each remaining trial, movement onset was defined as the time point at which this distance exceeded a threshold—five times the standard deviation of the resting signal (prior to the ‘go’ cue) —and remained above this threshold for at least 100 ms. RT was quantified as the interval between the ‘go’ cue and movement onset.

### EEG data analysis

Raw EEG data were segmented into 8-second epochs/trials, spanning from 4 seconds before to 4 seconds after the ‘uncertainty’ cue. Pre-processing was performed using the open-source FieldTrip Matlab toolbox, following the pipeline described by Herring et al. (2015) (**Supplementary Fig. 1, Oostenveld et al., 2011; Herring et al., 2015**), which included an Independent Component Analysis (ICA) for artefact removal. For rTMS conditions, EEG signals from channel C3 were used to automatically detect individual TMS pulses based on stimulation artefacts. Trials in which TMS pulses could not be identified were excluded. To mitigate the ‘ringing artefact’ caused by the hardware filter’s step response and subsequent software filtering, data from 5 ms before the first TMS pulse to 25 ms after the last TMS pulse were removed. The remaining time serious were concatenated and subjected to a FastICA to identify and remove components associated with exponential decay artefact, residual muscle artefact, eye-blinks, eye movements, line noise, and other non-TMS related muscle artefacts (**Ilmoniemi and Kičić, 2011; Jung et al., 2000; Korhonen et al., 2011**). On average, 5.67 ± 0.22 (SEM) out of 20 components were manually identified and removed. The cleaned components were then used to reconstruct the time serious, and the missing segment during TMS pulses were interpolated by mirroring the data segments immediately before and after the pulses to ensure the data continuity. For the no TMS condition, a similar preprocessing pipeline was applied, resulting in the removal of 5.41 ± 0.36 (SEM) out of 20 components.

The pre-processed EEG data were analysed using FieldTrip to investigate the associations between EEG power spectrum, uncertainty, motor control, and rTMS. Specifically, EEG signals were first band-pass filtered between 1-95 Hz, followed by a band-stop filter between 48-52 Hz, both implemented using 4^th^-order zero-phase Butterworth filters. The filtered signals were decomposed into time-frequency domain using continuous wavelet transformation with a linear frequency scale ranging from 1 to 95 Hz, at 1 Hz frequency resolution, and a wavelet width of 6 cycles. To derive the time course of power within specific frequency bands (e.g., IBF ± 3 Hz and alpha band), the power spectrum was averaged across selected frequencies and subsequently decibel (dB) normalised relative to the mean power in the 500 ms preceding the ‘uncertainty’ cue.

Furthermore, in order to investigate the potential effect of IBF phase when the first rTMS pulse was delivered, the phase at IBF of the pre-TMS EEG signal was estimated at channel C3 for each individual trial, following a similar procedure as used by van Elswijk et al. (**2010**) and Torrecillos et al. (**2020**). Briefly, an epoch lasting 2 cycles of the IBF ending prior the TMS pulse was extracted, multiplied by a Hanning taper, and then Fourier transformed to determine the signal’s phase. These initial phase values were then categorised into eight phase bins to assess potential phasic effects of rTMS (**Torrecillos et al. 2020; He et al., 2021**).

### Statistical analysis

Statistical analyses were performed using custom scripts written in MATLAB R2023b (The MathWorks Inc, Nantucket, MA).

To compare the group-averaged power time courses at different time points relative to the uncertainty or ‘go’ cue, a non-parametric cluster-based permutation test (1000 repetitions) was applied, following Maris and Oostenveld **(2007**). This method inherently controls the false alarm rate across multiple comparisons.

For trial-by-trial measures, including behavioural performance (RT), uncertainty levels (no, low, and high), stimulation conditions (no TMS, regular rTMS, and irregular rTMS), and beta band features (power and phase bins), generalised linear mixed effect (GLME) modelling was used to examine variable associations (**Yu et al., 2022; He et al., 2025**). In each GLME model, an independent random slope(s) between the predictor(s) and the dependent variable as well as an independent random intercept (s) were included. Parameters were estimated using maximum likelihood with Laplace approximation. Model outputs included the Akaike Information Criterion, coefficient estimates with standard errors (k ± SE), multiple comparison-corrected P-values, variable interactions (R2). False discovery rate (FDR) correction was applied to account for multiple comparisons across metrics (**Benjamini et al., 2001**). Full specifications of the GLME models are available in Supplementary Tables.

To assess beta phase–dependent effects of rTMS on RT, trials for each participant and rTMS condition were grouped into two bins. Specifically, trials with initial beta phases between –π and –π/4 were assigned to Phase 1 (“down state”), and those between 0 and 3π/4 to Phase 2 (“up state”) (**Salimpour et al., 2022; Guo et al., 2025**). GLME models were then used to examine the associations among phase bin, uncertainty level, and RT in both regular and irregular rTMS conditions.

## Data availability statement

The scripts and raw data required to reproduce the analyses in this article will be shared under a CC BY-SA license on the data-sharing platform of the MRC CoRE in Restorative Neural Dynamics.

## Results

### Movement initiation is modulated by both uncertainty and rTMS

To assess behavioural performance (RT) across different conditions, we first examined the main effects of uncertainty (no, low, and high) and rTMS (no, regular, and irregular), as well as their interaction, using a GLME model (**Supplementary Table 1, Model 1**). The analysis revealed that higher uncertainty was significantly associated with slower movement initiation, reflected in longer RT (k = 0.0680 ± 0.0042, P_FDR_ < 0.0001), as illustrated in **Fig. 2A**. A main effect of rTMS indicated that stimulation significantly shortened RT compared to no TMS condition (k = -0.0109 ± 0.0034, P_FDR_ = 0.0016). The interaction between uncertainty and rTMS was not significant (k = -0.0013 ± 0.0014, P = 0.3353), suggesting that the effect of ‘uncertainty’ during the ‘preparation phase’ was independent from the rTMS conditions. To further explore the effects of different rTMS protocols, we conducted separate GLME models, each comparing two of the three rTMS conditions (**Supplementary Table 1, Models 2-4**). These analysis showed that both regular (k = -0.0190 ± 0.0071, P_FDR_ = 0.0070) and irregular (k = -0.0107 ± 0.0036, P_FDR_ = 0.0028) rTMS significantly and comparably (k = -0.0034 ± 0.0053, P_FDR_ = 0.5242) reduced RT relative to the no TMS condition (**Fig. 2B**). These effects were consistent across different levels of uncertainty, as evidenced by the absence of significant interaction effects in all models.

**Fig. 2.**
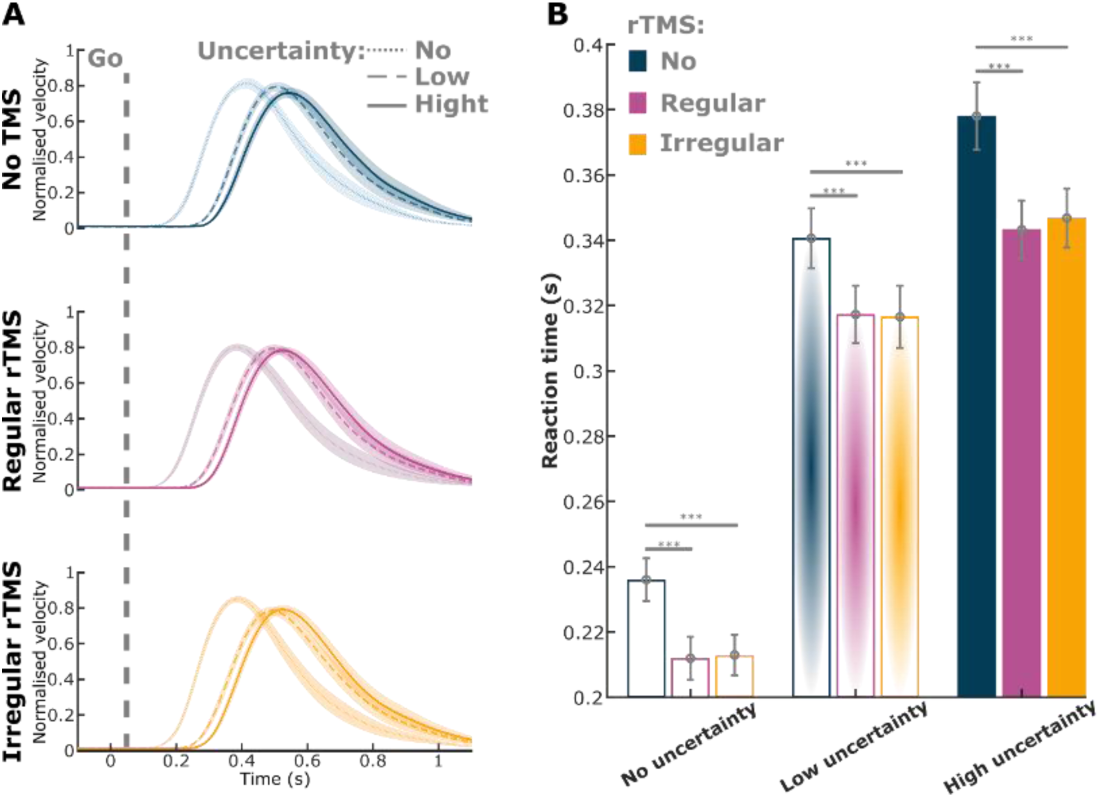
Behavioural results. **(A)** Normalised velocity time courses aligned to the ‘go’ cue (grey dashed line) are shown for different levels of uncertainty (solid, dotted, and dash lines) and rTMS conditions (green, pink, and yellow). **(B)** Reactions times are compared across uncertainty and rTMS conditions. Given the absence of a significant interaction between uncertainty and rTMS, the statistical results presented here reflect the main effects of rTMS on reaction time, as derived from the GLME models (**Supplementary Table 1**).

### Cortical beta and alpha are bilaterally and proportionally modulated by directional uncertainty

To investigate the modulatory effects over the primary motor cortices under varying levels of directional uncertainty, we analysed time-frequency power spectra from both hemispheres (electrodes C3 and C4), aligned to the onset of the ‘uncertainty’ cue, using the data recorded in the absence of rTMS. A clear bilateral reduction in beta-band power was observed following the ‘uncertainty’ cue (**Fig. 3A and D**), with the extent of modulation varying by uncertainty level: higher uncertainty was associated with a smaller reduction in beta power (i.e., higher absolute beta power). Beta activity further decreased after the ‘go’ cue and rebounded following movement onset (**Fig. 3B and E**). GLME modelling revealed significant main effects of uncertainty level (no, low, and high; k = 0.4108 ± 0.0696, P_FDR_ = 7.9494 × 10^-9^) and laterality (left vs. right; k = 0.4403 ± 0.1117, P_FDR_ = 8.2834 × 10^-5^) on beta power modulation during the ‘preparation phase’, suggesting stronger beta modulation in the left hemisphere (contralateral to the task hand) under lower uncertainty, as also illustrated in the topographic maps (**Fig. 3G**). However, no significant interaction was found between uncertainty level and laterality. Pairwise comparisons revealed significant differences in beta modulation between the no and high uncertainty conditions, as well as between low and high uncertainty, in both hemispheres (**Fig. 3C and F**). Similar patterns were also observed in the alpha frequency band (**Supplementary Fig. 2**). Interestingly, although the left hemisphere exhibited greater overall beta and alpha modulation following the ‘uncertainty’ cue, GLME modelling revealed that modulation in the right hemisphere more accurately predicted the associated level of uncertainty. This was further supported by **Fig. 3H**, which shows a broader area of larger beta modulation difference between high and no uncertainty in the right hemisphere. Together, these results suggest that both beta and alpha activity are bilaterally modulated by directional uncertainty during movement preparation, with stronger modulation in the left hemisphere and more precise representation of uncertainty levels in the right. Details of GLME models can be found in **Supplementary Table 2**.

**Fig. 3.**
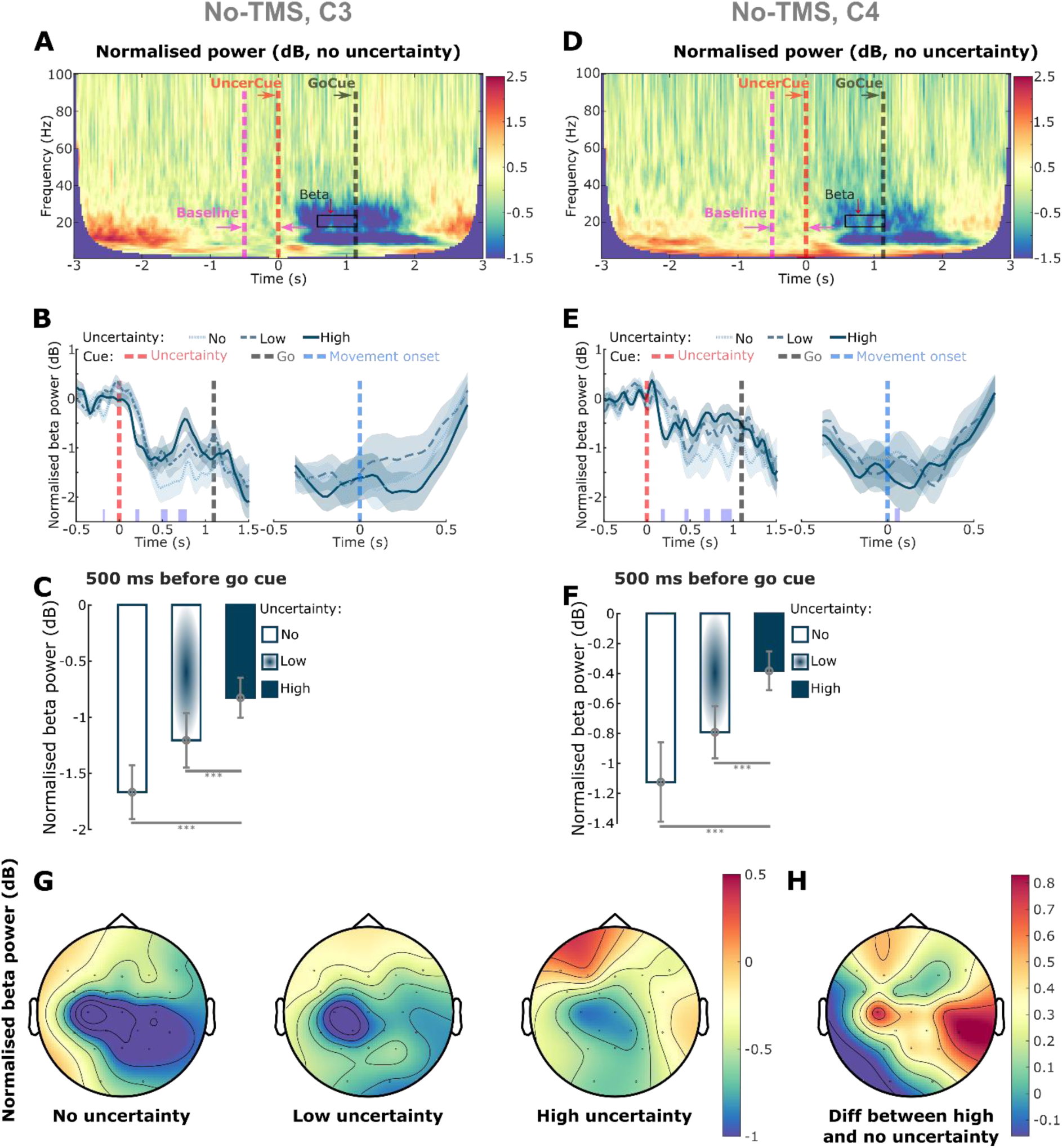
Bilateral modulation of beta activity by uncertainty. **(A)** Group-averaged time-frequency power spectra from EEG channel C3 (left hemisphere), aligned to the onset of the ‘uncertainty’ cue (red dashed line), in the no-rTMS/no-uncertainty condition. Power spectra were normalised to a 500-millisecond pre-cue resting baseline for each trial. Beta-band power (black box) showed modulations following the ‘uncertainty’ cue and prior to the ‘go’ cue (black dashed line). **(B)** Group-averaged time courses of beta power, normalised to baseline, in different uncertainty conditions. Red, black, and blue dashed lines indicate the ‘uncertainty’, ‘go’, and ‘movement onset’ cues, respectively. **(C)** Comparison of beta power modulation across different uncertainty conditions. Power was quantified as the average within 500-millisecond window preceding the ‘go’ cue, normalised (in dB) to the 500-millisecond pre-‘uncertainty’ cue baseline. Error bars represent the mean ± SEM across participants. **(D)-(F)** Same analyses as in (A)-(C), but for EEG channel C4 (right hemisphere). **(G)** Topographical map of beta power modulation under conditions of no (left), low (middle), and high (right) uncertainty. Power was quantified as in (C) and (F). **(H)** Difference in beta power modulation between high and no uncertainty conditions. *P < 0.05, **P < 0.01, ***P < 0.001. P-values in (C) and (F) were derived from generalised linear mixed-effects models applied to individual trials and corrected for multiple comparisons using FDR. Purple bars in (B) and (E) indicate significant differences between no- and high-uncertainty conditions (cluster-based permutation test).

### Movement initiation is predicted by contralateral cortical beta preceding the ‘go’ cue

We demonstrated that directional uncertainty modulates both behavioural performance (e.g., RT) and bilateral cortical beta/alpha activity. To investigate the relationship between these neural and behavioural modulations across varying levels of uncertainty, we applied GLME models to predict RT based on beta/alpha power immediately preceding (500 ms) the ‘go’ cue, while controlling for baseline differences across participants and uncertainty conditions. The results of this analysis (**Table 1**) revealed that, after considering the effect of uncertainty level, higher contralateral beta power before the ‘go’ cue was significantly associated with longer RT (k = 0.0017 ± 0.0008, P_FDR_ = 0.0327). In contrast, contralateral alpha (k = -0.0011 ± 0.0006, P_FDR_ = 0.0781), ipsilateral beta (k = 0.0010 ± 0.0007, P_FDR_ = 0.3048), and ipsilateral alpha (k = 0.0002 ± 0.0006, P_FDR_ = 0.7348) did not significantly predict RT, although all were modulated by uncertainty.

**Table 1.**
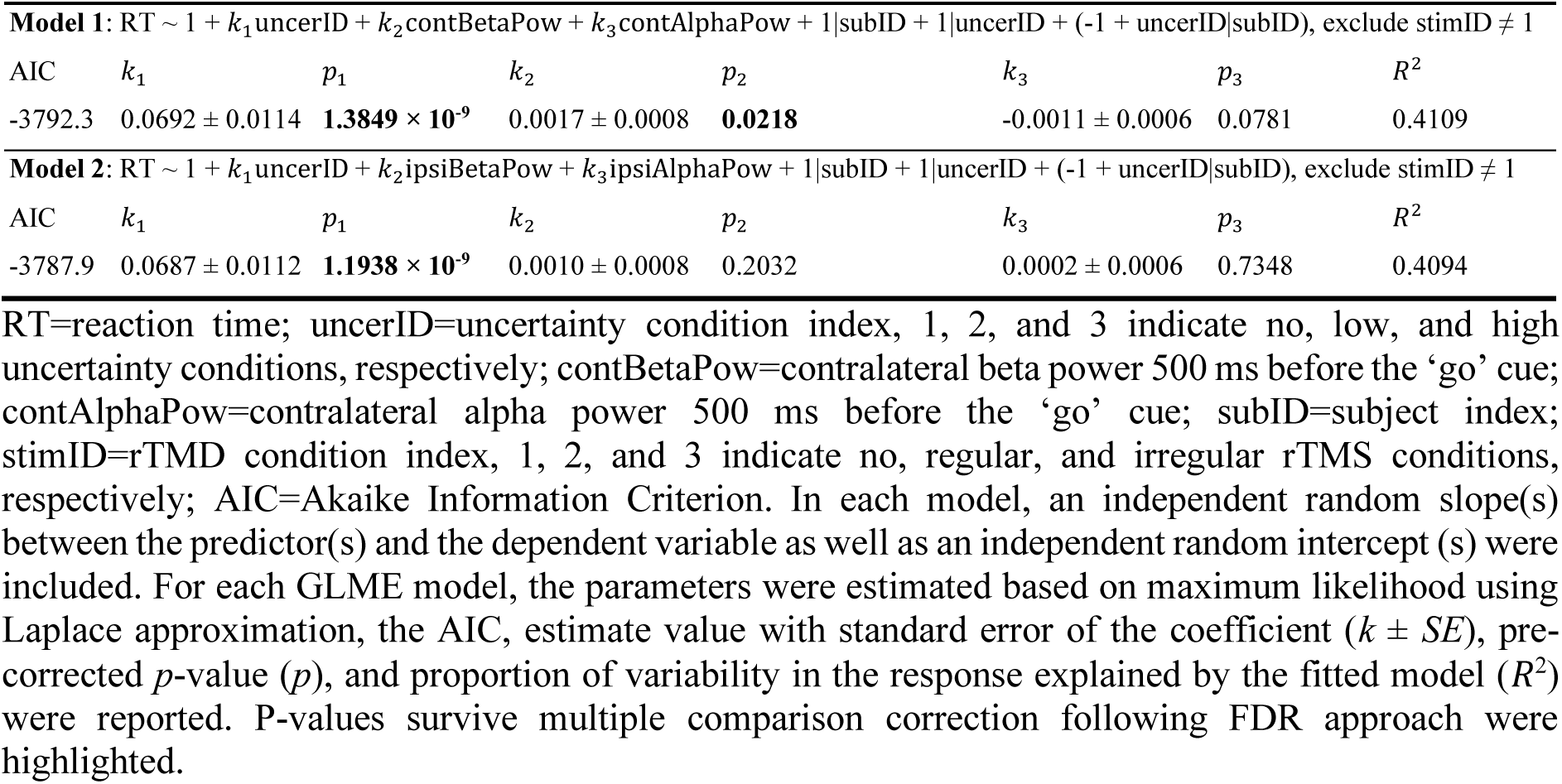
Prediction of RT based on cortical beta and alpha power preceding the ‘go’ cue using generalised linear mixed effect (GLME) modelling.

| **Model 1:** $RT \sim 1 + k_1 \text{uncerID} + k_2 \text{contBetaPow} + k_3 \text{contAlphaPow} + 1\vert \text{subID} + 1\vert \text{uncerID} + (-1 + \text{uncerID}\vert \text{subID})$, exclude $\text{stimID} \neq 1$ | | | | | | | |
| --- | --- | --- | --- | --- | --- | --- | --- |
| AIC | $k_1$ | $p_1$ | $k_2$ | $p_2$ | $k_3$ | $p_3$ | $R^2$ |
| -3792.3 | $0.0692 \pm 0.0114$ | **$1.3849 \times 10^{-9}$** | $0.0017 \pm 0.0008$ | **0.0218** | $-0.0011 \pm 0.0006$ | 0.0781 | 0.4109 |
| **Model 2:** $RT \sim 1 + k_1 \text{uncerID} + k_2 \text{ipsiBetaPow} + k_3 \text{ipsiAlphaPow} + 1\vert \text{subID} + 1\vert \text{uncerID} + (-1 + \text{uncerID}\vert \text{subID})$, exclude $\text{stimID} \neq 1$ | | | | | | | |
| AIC | $k_1$ | $p_1$ | $k_2$ | $p_2$ | $k_3$ | $p_3$ | $R^2$ |
| -3787.9 | $0.0687 \pm 0.0112$ | **$1.1938 \times 10^{-9}$** | $0.0010 \pm 0.0008$ | 0.2032 | $0.0002 \pm 0.0006$ | 0.7348 | 0.4094 |

RT=reaction time; uncerID=uncertainty condition index, 1, 2, and 3 indicate no, low, and high uncertainty conditions, respectively; contBetaPow=contralateral beta power 500 ms before the ‘go’ cue; contAlphaPow=contralateral alpha power 500 ms before the ‘go’ cue; subID=subject index; stimID=rTMD condition index, 1, 2, and 3 indicate no, regular, and irregular rTMS conditions, respectively; AIC=Akaike Information Criterion. In each model, an independent random slope(s) between the predictor(s) and the dependent variable as well as an independent random intercept (s) were included. For each GLME model, the parameters were estimated based on maximum likelihood using Laplace approximation, the AIC, estimate value with standard error of the coefficient ( $k \pm SE$ ), pre-corrected $p$ -value ( $p$ ), and proportion of variability in the response explained by the fitted model ( $R^2$ ) were reported. P-values survive multiple comparison correction following FDR approach were highlighted.

### rTMS facilitates movement initiation through attenuating beta ERD

We found that both regular and irregular rTMS significantly shortened RT compared to the no TMS condition (**Fig. 2**), with no significant difference between the two rTMS conditions. GLME modelling indicated that the reduction in RT under rTMS conditions could not be explained by the timing of the first/last TMS pulse (**Supplementary Table 3**), suggesting that the observed behavioural effects are unlikely to be attributable to placebo timing influences of the auditory or tactile sensations associated with the rTMS pulses. To investigate the effects of rTMS on beta activity, we quantified beta event-related desynchronisation (ERD) around movement initialisation by calculating the difference in beta power between a baseline period (500 ms prior to the ‘uncertainty’ cue) and a 500-ms window centred around movement onset. We then compared beta ERD across different uncertainty and stimulation conditions using GLME modelling. As illustrated in **Fig. 4A**, beta ERD was not significantly modulated by uncertainty during movement preparation in all three stimulation conditions. However, rTMS significantly reduced beta ERD compared to the no TMS condition (No TMS vs. Regular rTMS: k = - 0.6279 ± 0.2241, P_FDR_ = 0.0231; No TMS vs. Irregular rTMS: k = -0.3003 ± 0.1126, P_FDR_ = 0.0231), with no significant difference observed between the two rTMS conditions (k = 0.0260 ± 0.0885, P_FDR_ = 0.8454). It is worth noting that the average frequency of the irregular TMS condition is also the IBF, as the timing of the last pulse with respect to the first pulse matched those of the IBF rTMS condition. This modulation effect was specific to the stimulation site (C3) at the beta frequency band and was not observed at the other site (C4, **Fig. 4B**) or within the alpha frequency band (**Supplementary Fig. 3**). To further investigate the relationship between the behavioural and neuronal effects of rTMS, we first quantified baseline RT and beta ERD around movement initialisation by averaging values across all no-TMS trials within each uncertainty condition. Using these baselines, we then calculated trial-wise reductions in RT and beta ERD for each rTMS trial. Here regular and irregular rTMS conditions were pooled together, as no significant differences were observed between them in either behavioural or neural measures. We applied a GLME model to assess the relationship between changes in beta ERD around movement initialisation and RT under rTMS. The analysis revealed that greater reductions in beta ERD, i.e. less beta reduction around movement initialisation, significantly predicted grater reductions in RT (k = 0.0006 ± 0.0003, P = 0.0364); however, this effect did not survive correction for multiple comparisons (P_FDR_ = 0.0727). Notably, the effect was stronger and remained significant when using a broad beta frequency band (13-30 Hz) instead of the IBF (k = 0.0010 ± 0.0004, P_FDR_ = 0.0082) but was not significant when beta was replaced with alpha (k = 0.0004 ± 0.0002, P_FDR_ = 0.1177).

**Fig. 4.**
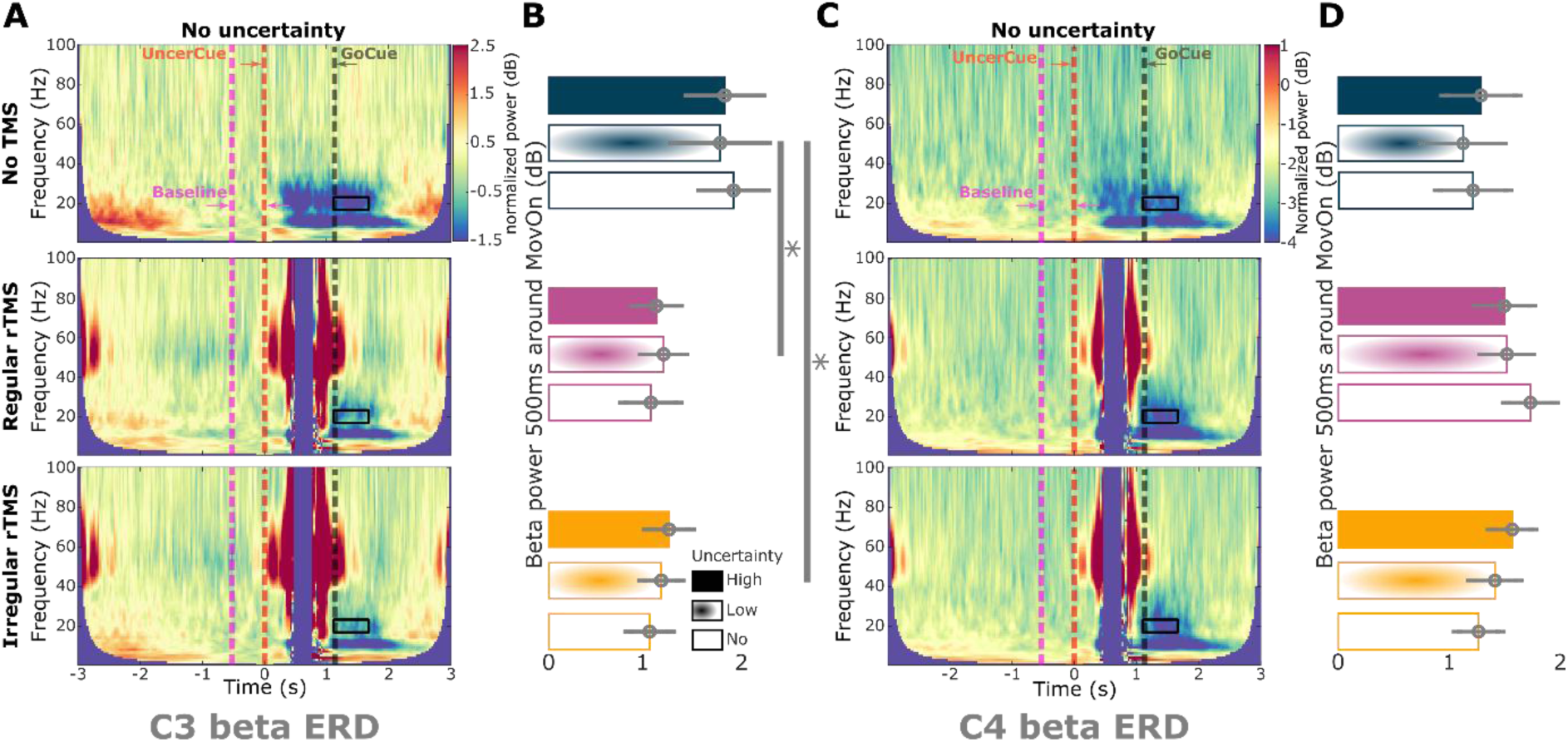
Unilateral modulation of beta event-related desynchronisation by rTMS. **(A)** Group-averaged time-frequency power spectra from EEG channel C3 (left hemisphere), aligned to the onset of the ‘uncertainty’ cue (red dashed line), shown for three conditions: no uncertainty and no TMS (up), regular rTMS (middle), and irregular rTMS (bottom). Power spectra were normalised to a 500-millisecond pre-cue resting baseline (pink dashed line) for each trial. Beta-band activity (black box) showed modulation following movement onset (black dashed line). **(B)** Comparison of beta ERD across different uncertainty and rTMS conditions. Power was quantified as the average within a 500-ms window preceding the ‘uncertainty’ cue and normalised (in dB) to the 500 ms window centered at movement onset. Error bars represent mean ± SEM across participants. **(C)-(D)** Same analyses as in (A)-(B), but for EEG channel C4 (right hemisphere). *P < 0.05. P-values in (B) and (D) were obtained using generalised linear mixed-effects models applied to individual trials and corrected for multiple comparisons using FDR.

### The effects of rTMS on movement initiation depends on the initial phase of the targeted beta activity

To further investigate the potential effects of the brain state when the rTMS started, we performed a beta phase-dependent analysis (**Fig. 5A**, see Methods). In the regular rTMS condition, RTs were significantly shorter when rTMS started in Phase 1 (down phase) than in Phase 2 (up phase) of beta (k = 0.0079 ± 0.0032, P_FDR_ = 0.0472; **Fig. 5B**), an effect that could not be explained by differences in uncertainty (k = 0.0244 ± 0.0386, P_FDR_ = 0.5993; **Fig. 5C**) or by pre-stimulation beta power (k = -0.0941 ± 0.1792, P_FDR_ = 0.5993; **Fig. 5D**). In contrast, we did not find a statistically significant phase-dependent modulation of RT for the irregular rTMS condition (p > 0.05) (**Fig. 5E-G**). Moreover, RTs when regular rTMS started in Phase 1 were significantly shorter than those in the irregular rTMS condition (k = 0.0027 ± 0.0013, P = 0.0352).

**Fig. 5.**
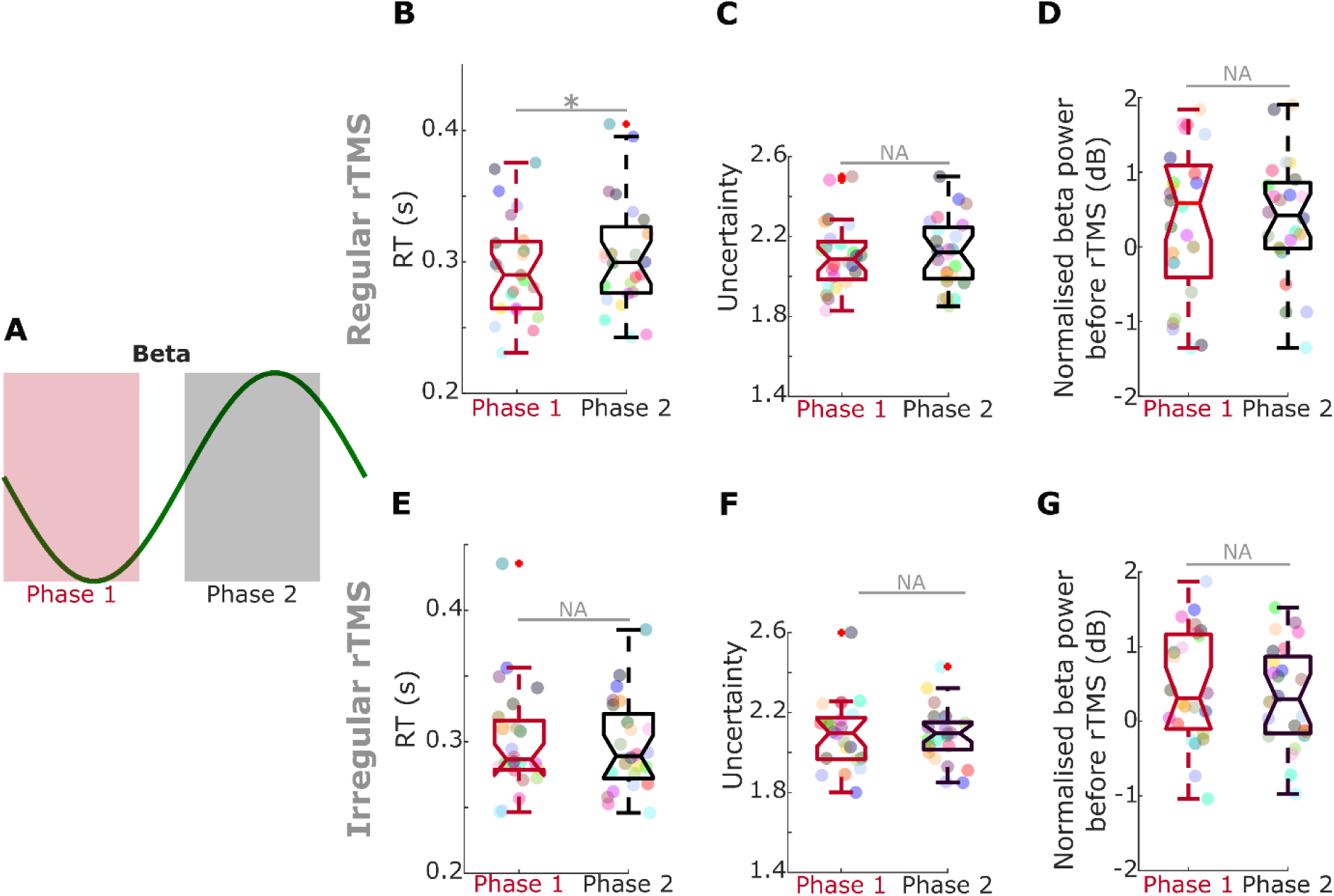
Phasic effects of regular and irregular rTMS. (A) A. **s**chematic illustration showing how two-phase bins (‘down state’ and ‘up state’) are defined. **(B)-(D)** Comparisons between the two-phase groups in terms of RTs (B), uncertainty levels (C), and beta power preceding stimulation (D). Error bars indicate mean ± SEM across participants. *P < 0.05. P-values were derived from generalised linear mixed-effects models applied to individual trials and corrected for multiple comparisons using FDR. **(E)-(G)** Same analyses as in (B)-(D), but under the irregular rTMS condition.

## Discussion

Beta-band oscillations in the cortical-basal ganglia network have been linked to both physiological motor (**Little et al., 2019; He et al., 2020a; Wessel, 2020**) and cognitive (**Tzagarakis et al., 2010; Tan et al., 2016**) functions, as well as pathological conditions such as Parkinson’s disease (**Jenkinson and Brown, 2011; Crowell et al., 2012; Cagnan et al., 2019**). However, the extent to which cognition-versus motor-related beta oscillations differ, and how we can better modulate them, remains incompletely understood. In this study, we combined a cognitive-motor behavioural task and non-invasive rTMS intervention to investigate the distinct roles of cortical beta oscillations in relation to uncertainty, motor control, and their modulation with rTMS.

### Beta-band oscillations during movement preparation encode uncertainty

Previous studies have shown that beta activity in the sensorimotor cortex scales with directional uncertainty during movement preparation (**Tzagarakis et al., 2010**), beta/alpha band power over central cortical regions decreases more in conditions with predetermined/predictable movements (i.e., no uncertainty) compared to conditions requiring a choice/non-predictable (**Manuel et al., 2003; Helvert et al., 2021**). Consistent with these previous findings, in this study, we found that cortical beta and alpha in both the left and right hemispheres decreased during movement preparations following the ‘uncertainty’ cue, with a smaller reduction (i.e., higher beta) observed under higher uncertainty condition (**Fig. 3 and Supplementary Fig. 2**). This effect may be attributed to the suggested associations between inhibitory interneurons and beta-band oscillations (**Jensen et al., 2005; Yamawaki et al., 2008**). Specifically, increased inhibition under high uncertainty conditions and decreased inhibition under low uncertainty conditions may underlie the observed differences in beta-band activity modulation. Furthermore, even though we found that overall modulation of beta and alpha activity was greater in the left hemisphere (contralateral to the upcoming movement), beta modulations in the right hemisphere more closely tracked uncertainty levels. This observation aligns with findings from **Tzagarakis et al. (2010)**, who reported that a greater number of channels in the left hemisphere exhibited pronounced beta-band power decreases during the peri-movement period under a similar paradigm. It has been suggested that the left hemisphere is primarily involved in processing temporal features such as duration, sequencing, and rhythm, while the right hemisphere is more engaged in spatial mapping and body positioning (**Bradshaw & Nettleton, 1981**). In the context of uncertainty, the left hemisphere has been associated with generating explanations and filling informational gaps, whereas the right hemisphere contributes to conflict detection, belief updating, and cognitive flexibility (**Marinsek et al., 2014**).

### Beta-band oscillations before movement encode reaction time

Modulation of cortical beta-band oscillations by movement has been well established over the past decades (**Pfurtscheller et al., 1996; Pfurtscheller and Silva, 1999; Jurkiewicz et al., 2006**). Specifically, EEG power is known to decrease within the alpha-beta (or ‘mu’) rhythm during movement preparation—known as event-related desynchronisation (ERD), and to increase following movement initiation—referred to as event-related synchronisation (ERS). Our data similarly demonstrated clear beta ERD preceding movement onset and beta ERS following it (**Fig. 3 and Supplementary Fig. 2**). Given that higher uncertainty conditions were associated with significantly longer RTs (Fig. 2) and elevated beta power, we accounted for baseline differences in beta activity and RT across uncertainty levels. The results showed that greater beta power (but not alpha) prior to the ‘go’ cue was associated with slower movement initiation (i.e., longer RTs) (**Table 1**). This finding aligns with previous research showing that elevated beta activity in the cortical-basal ganglia network is predictive of delayed movement initiation, both in healthy individuals (**Tzagarakis et al., 2010; Little et al., 2019; Wessel et al., 2020; He et al., 2020a**) and in patients with Parkinson’s disease (**Torrecillos et al., 2018; Lofredi et al., 2019; He et al., 2020b**).

Despite significant differences in behavioural performance and beta oscillatory dynamics across uncertainty conditions, beta power consistently decreased to a similar minimal level around movement onset, a phenomenon also reported by **Tzagarakis et al. (2010)**. Concretely, trials with slower movement initiation—whether due to higher uncertainty or elevated pre-movement beta—were associated with a larger reduction in beta power, converging toward this minimal level. Indeed, it has been found that during self-paced movements, corticospinal excitability increases and reaches a maximal level during movement initiation, then reduces after movement initiation, presenting a very similar pattern to beta ERD (**Chen et al., 1998**). Based on these observations, we speculate that the low beta power observed at movement onset may reflect a neural threshold, marked by beta ERD, that must be reached to successfully initiate movement. This speculation is further supported by findings on the effects of normal aging on beta ERD. Specifically, older adults exhibit greater beta ERD amplitudes over regions contralateral to the active effector compared to younger individuals across a variety of motor tasks, as shown in both EEG and MEG studies (**Sailer et al., 2000; Mary et al., 2015; Toledo et al., 2016; Johari and Behroozmand, 2020; Walker et al., 2020; Peter et al., 2022**). Additionally, elite athletes have been shown to exhibit globally lower beta ERD amplitudes than non-athletes while performing the same air-pistol shooting task (**Del Percio et al., 2009**).

### Effects of regular and irregular rTMS

Numerous studies report that high-frequency rTMS (≥ 5 Hz) generally increases cortical excitability and is associated with reduced RTs (**Pascual-Leone et al., 1998; Huang et al., 2005**; **Vanderhasselt et al., 2006; Kim et al., 2012; Spagnolo et al., 2014; Li et al., 2017; Parris et al., 2021; Asl and Vaghef, 2022**), whereas low-frequency rTMS (< 5 Hz) tends to decrease excitability and is linked to prolonged RTs (**Chen et al., 1997**; **Pascual-Leone et al., 1998; Schlaghecken et al., 2003; Terao et al., 2007; Brusa et al., 2009; Peterchev et al., 2012; Spagnolo and Goldman, 2017**). Consistent with this literature, we observed that RTs were significantly reduced under rTMS relative to the no-TMS condition across all uncertainty levels (**Fig. 2**). In parallel, beta-band ERD—but not alpha-band ERD—was significantly smaller under rTMS than in the no-TMS condition (**Fig. 4 and Supplementary Fig. 3**). Importantly, the reduction in beta ERD (but not alpha) significantly predicted reductions in reaction times produced by rTMS across uncertainty levels. Together, these results suggest that high frequency rTMS may facilitate movement initiation by increasing cortical excitability, resulting in a lower threshold for movement initiation that represented as a smaller beta ERD around movement initialisation.

However, we did not observe significant differences in either RT or beta ERD between the regular and irregular rTMS conditions. This lack of differentiation may be attributed to the following three factors. First, in addition to frequency, other stimulation parameters—number of pulses, stimulation intensity, duration, inter-train interval, and coil position—are known to influence the degree of excitability modulation (**Huang et al., 2005; Pell et al., 2011; Rossi et al., 2021**). In this study, we carefully matched all stimulation parameters between the two rTMS conditions except for frequency. As a result, the measured outcomes (i.e., RT and beta ERD) may not have been sensitive enough to detect frequency-specific effects of rTMS. Second, the instantaneous frequencies between TMS pulses were relatively high in both rTMS conditions—18.96-23.01 Hz (5^th^–95^th^ percentile) in the regular condition and 12.49-49.95 Hz (5^th^–95^th^ percentile) in the irregular condition (**Supplementary Fig. 4**). These high frequencies may have elicited comparable excitatory effects on cortical activity (**Pascual-Leone et al., 1998; Huang et al., 2005**; **Vanderhasselt et al., 2006; Kim et al., 2012; Spagnolo et al., 2014; Li et al., 2017; Parris et al., 2021; Asl and Vaghef, 2022**). Third, the efficacy of rTMS is also believed to depend on the brain state at the time of stimulation, including the phase of the ongoing oscillations (**Mitchell et al., 2007; Pell et al., 2011; Zanos et al., 2018; Zrenner et al., 2018; Schaworonkow et al., 2019; Torrecillos et al., 2020**). In line with this, we observed a significant phase-dependent reduction in RTs in the regular rTMS condition, but not in the irregular condition (**Fig. 4**). Specifically, rTMS delivered at the ‘down state’ of endogenous beta oscillation had a strongest effect in reducing RT compared to regular rTMS delivered to the ‘up state’ of beta and irregular rTMS. This is consistent with previous studies showing that TMS applied to the primary motor cortex (M1) targeting the sensorimotor mu-rhythm trough phase has been associated with higher corticospinal excitability than during the peak phase, as measured by the amplitude of motor evoked potentials (MEP) (**Zrenner et al., 2018; Hougland et al., 2025**). Regular rTMS starting at the ‘down state’ of endogenous beta oscillations has more likelihood to target the same phase over multiple consecutive beta cycles compared to irregular rTMS, which main explain the phase dependent effect of regular rTMS. This also suggest that continuous cycle-by-cycle rTMS pulses phase-locked to specific beta phases may be most effective in modulating cortical excitability and movement initialisation.

### Implications in Parkinson’s disease

Although older adults generally exhibit greater beta ERD compared to younger individuals (**Sailer et al., 2000; Mary et al., 2015; Toledo et al., 2016; Johari and Behroozmand, 2020; Walker et al., 2020; Peter et al., 2022**), reduced beta ERD has been reported in people with PD compared to age-matched healthy controls (**Heinrichs-Graham et al., 2014**). This attenuation may be attributed to elevated baseline beta power, a hallmark of PD pathophysiology (**Kühn et al., 2006; Brittain and Brown, 2014; Oswal et al., 2016**), which diminishes the relative change in beta amplitude during movement initiation, i.e., the ERD. Both L-Dopa and DBS have been shown to enhance beta ERD compared to the OFF-therapy state (**Doyle et al., 2005; Mathiopoulou et al., 2024**). These effects are likely mediated through the suppression of pathological beta oscillations within the cortical-basal ganglia network (**Kühn et al., 2009; Neumann et al., 2016; Kehnemouyi et al., 2021**). Recent evidence suggests that both medication and STN DBS may exert their therapeutic effects via a shared mechanism: disruption of high-beta band hyperdirect pathway coupling between cortex and basal ganglia, which is associated with excessive inhibitory drive (**Oswal et al., 2021; Binns et al., 2025**). In addition to medication and DBS, various non-invasive neuromodulation approaches—including rTMS, transcranial direct current stimulation (tDCS), temporal interference stimulation (TIS), and low-intensity transcranial focused ultrasound stimulation (LItFUS)—have been explored for PD treatment (**Chou et al., 2015; Duan and Zhang, 2024; Lamoš et al., 2025; Darmani et al., 2025**). However, their stimulation parameters, targets, clinical efficacy, and underlying mechanisms remain inconsistent and less well understood. The findings of this study may help clarify some of these discrepancies. For instance, rTMS effects in PD appear more robust when high-frequency stimulation is applied over M1, or when low-frequency stimulation targets frontal regions (**Chou et al., 2015**). These outcomes may reflect the excitatory effects of high-frequency rTMS on motor cortex, as suggested in this study, and the modulation of the abnormal prefrontal-subthalamic hyperdirect pathway implicated in PD (**Chen et al., 2020**).

## Limitations

There are several limitations in the current study. First, EEG recordings were conducted using a relatively low-density setup with only 23 channels. As a result, motor cortical activity was interpreted at the sensor level, which likely reflects cortical field potentials rather than direct neural sources. Future studies could enhance spatial resolution by employing high-density EEG systems. Second, cortical excitability under the two rTMS conditions was inferred from reaction times and beta ERDs, since rTMS was delivered subthreshold and did not elicit MEPs that could be used for direct excitability measurement. Third, the potential phasic effects of rTMS were identified through post hoc analysis, as the study was not originally designed to investigate phase-specific modulation. Emerging methodologies now enable precise cycle-by-cycle phase-locked stimulation (e.g., **McNamara et al., 2022; Guo et al., 2025**), which could be leveraged in future research to more rigorously test the phasic effects of IBF rTMS observed in this study.

## Conclusion

This study demonstrates that cortical beta oscillations are involved in the encoding of both directional uncertainty and movement preparation and execution, and that these cortical oscillations can be modulated by high frequency rTMS. Presentation of ‘uncertainty’ cues led to bilaterally suppression of beta power, with greater uncertainty associated with a smaller reduction in beta activity. In contrast, movement-related beta modulation was primarily lateralised to the hemisphere contralateral to the executing hand, where elevated beta power prior to the ‘go’ cue was associated with delayed movement initiation, as reflected in longer RTs. Importantly, high frequency rTMS significantly shortened RTs and attenuated beta ERD compared to the no TMS condition. The degree of beta ERD reduction correlated with the improvement in RTs, suggesting that beta ERD during movement initiation may reflect a neural threshold that must be overcome to initiate movement. While this threshold appears to be unaffected by uncertainty, it can be lowered through rTMS, thereby facilitating faster movement initiations.

## Acknowledgement

We thank all the participants for making this study possible. This work was supported by the Medical Research Council (MC_UU_00003/2) and the Guarantors of Brain. S.H. was also supported by The Royal Society (IES\R3\213123) and Parkinson’s UK.

---

**Supplementary Table 1.** Comparisons on reaction times across different uncertainty and rTMS conditions using generalised linear mixed effect (GLME) modelling.

|  |  |  |  |  |  |  |  |
| --- | --- | --- | --- | --- | --- | --- | --- |
| **Model 1:** $RT \sim 1 + k_1 \text{uncerID} + k_2 \text{stimID} + k_{inter} \text{uncerID} * \text{stimID} + 1\vert \text{subID} + (-1 + \text{uncerID}\vert \text{subID}) + (-1 + \text{stimID}\vert \text{subID})$ | | | | | | | |
| AIC | $k_1$ | $p_1$ | $k_2$ | $p_2$ | $k_{inter}$ | $p_{inter}$ | $R^2$ |
| -15611 | $0.0680 \pm 0.0042$ | < **0.0001** | $-0.0109 \pm 0.0034$ | **0.0016** | $-0.0013 \pm 0.0014$ | 0.3353 | 0.4565 |
| **Model 2:** $RT \sim 1 + k_1 \text{uncerID} + k_2 \text{stimID} + k_{inter} \text{uncerID} * \text{stimID} + 1\vert \text{subID} + (-1 + \text{uncerID}\vert \text{subID}) + (-1 + \text{stimID}\vert \text{subID})$, exclude $\text{stimID} = 3$ | | | | | | | |
| AIC | $k_1$ | $p_1$ | $k_2$ | $p_2$ | $k_{inter}$ | $p_{inter}$ | $R^2$ |
| -9639.5 | $0.0742 \pm 0.0055$ | < **0.0001** | $-0.0190 \pm 0.0071$ | **0.0070** | $-0.0055 \pm 0.0030$ | 0.0621 | 0.4402 |
| **Model 3:** $RT \sim 1 + k_1 \text{uncerID} + k_2 \text{stimID} + k_{inter} \text{uncerID} * \text{stimID} + 1\vert \text{subID} + (-1 + \text{uncerID}\vert \text{subID}) + (-1 + \text{stimID}\vert \text{subID})$, exclude $\text{stimID} = 2$ | | | | | | | |
| AIC | $k_1$ | $p_1$ | $k_2$ | $p_2$ | $k_{inter}$ | $p_{inter}$ | $R^2$ |
| -9660.5 | $0.0707 \pm 0.0047$ | < **0.0001** | $-0.0107 \pm 0.0036$ | **0.0028** | $-0.0019 \pm 0.0015$ | 0.2016 | 0.4530 |
| **Model 4:** $RT \sim 1 + k_1 \text{uncerID} + k_2 \text{stimID} + k_{inter} \text{uncerID} * \text{stimID} + 1\vert \text{subID} + (-1 + \text{uncerID}\vert \text{subID}) + (-1 + \text{stimID}\vert \text{subID})$, exclude $\text{stimID} = 1$ | | | | | | | |
| AIC | $k_1$ | $p_1$ | $k_2$ | $p_2$ | $k_{inter}$ | $p_{inter}$ | $R^2$ |
| -12104 | $0.0586 \pm 0.0067$ | < **0.0001** | $-0.0034 \pm 0.0053$ | 0.5242 | $0.0022 \pm 0.0024$ | 0.3649 | 0.4987 |

RT=reaction time; uncerID=uncertainty condition index, 1, 2, and 3 indicate no, low, and high uncertainty conditions, respectively; stimID=rTMD condition index, 1, 2, and 3 indicate no, regular, and irregular rTMS conditions, respectively; inter=interaction; subID=subject index; AIC=Akaike information criterion. In each model, an independent random slope(s) between the predictor(s) and the dependent variable as well as an independent random intercept (s) were included. For each GLME model, the parameters were estimated based on maximum likelihood using Laplace approximation, the AIC, estimate value with standard error of the coefficient ( $k \pm SE$ ), pre-corrected $p$ -value ( $p$ ), and proportion of variability in the response explained by the fitted model ( $R^2$ ) were reported. P-values survive multiple comparison correction following FDR approach were highlighted.

**Supplementary Table 2.**
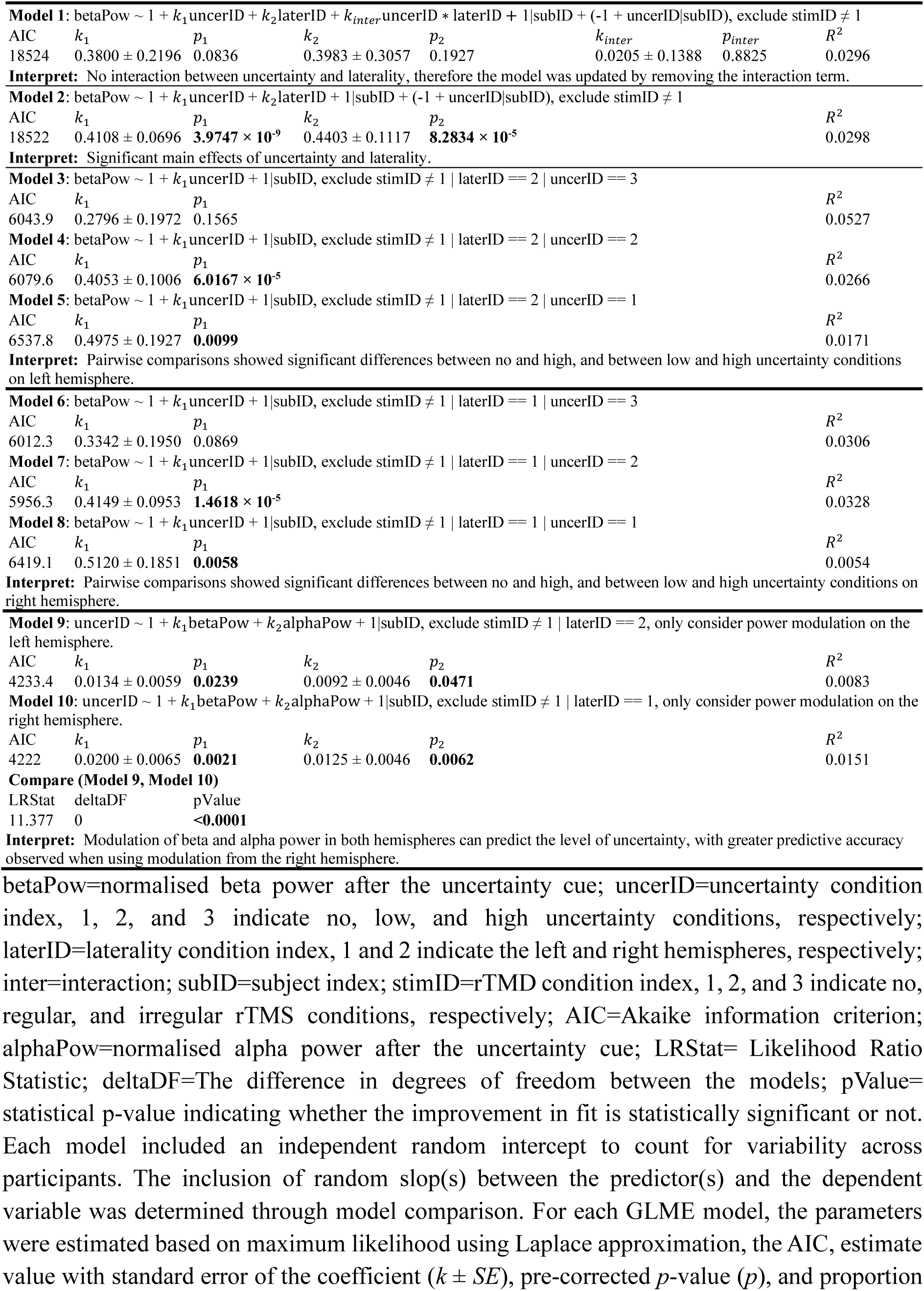
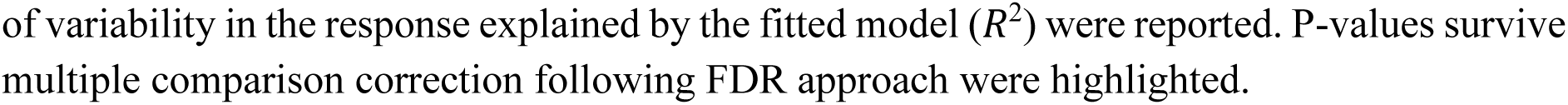
Comparisons on bilateral beta modulation across different uncertainty conditions in the absence of rTMS using generalised linear mixed effect (GLME) modelling.

|  |  |  |  |  |  |  |  |
| --- | --- | --- | --- | --- | --- | --- | --- |
| **Model 1:** betaPow ~ 1 + k<sub>1</sub>uncerID + k<sub>2</sub>laterID + k<sub>inter</sub>uncerID * laterID + 1\|subID + (-1 + uncerID\|subID), exclude stimID ≠ 1 |  |  |  |  |  |  |  |
| AIC | k<sub>1</sub> | p<sub>1</sub> | k<sub>2</sub> | p<sub>2</sub> | k<sub>inter</sub> | p<sub>inter</sub> | R<sup>2</sup> |
| 18524 | 0.3800 ± 0.2196 | 0.0836 | 0.3983 ± 0.3057 | 0.1927 | 0.0205 ± 0.1388 | 0.8825 | 0.0296 |
| **Interpret:** No interaction between uncertainty and laterality, therefore the model was updated by removing the interaction term. |  |  |  |  |  |  |  |
| **Model 2:** betaPow ~ 1 + k<sub>1</sub>uncerID + k<sub>2</sub>laterID + 1\|subID + (-1 + uncerID\|subID), exclude stimID ≠ 1 |  |  |  |  |  |  |  |
| AIC | k<sub>1</sub> | p<sub>1</sub> | k<sub>2</sub> | p<sub>2</sub> |  |  | R<sup>2</sup> |
| 18522 | 0.4108 ± 0.0696 | **3.9747 × 10<sup>-9</sup>** | 0.4403 ± 0.1117 | **8.2834 × 10<sup>-5</sup>** |  |  | 0.0298 |
| **Interpret:** Significant main effects of uncertainty and laterality. |  |  |  |  |  |  |  |
| **Model 3:** betaPow ~ 1 + k<sub>1</sub>uncerID + 1\|subID, exclude stimID ≠ 1 \| laterID == 2 \| uncerID == 3 |  |  |  |  |  |  |  |
| AIC | k<sub>1</sub> | p<sub>1</sub> |  |  |  |  | R<sup>2</sup> |
| 6043.9 | 0.2796 ± 0.1972 | 0.1565 |  |  |  |  | 0.0527 |
| **Model 4:** betaPow ~ 1 + k<sub>1</sub>uncerID + 1\|subID, exclude stimID ≠ 1 \| laterID == 2 \| uncerID == 2 |  |  |  |  |  |  |  |
| AIC | k<sub>1</sub> | p<sub>1</sub> |  |  |  |  | R<sup>2</sup> |
| 6079.6 | 0.4053 ± 0.1006 | **6.0167 × 10<sup>-5</sup>** |  |  |  |  | 0.0266 |
| **Model 5:** betaPow ~ 1 + k<sub>1</sub>uncerID + 1\|subID, exclude stimID ≠ 1 \| laterID == 2 \| uncerID == 1 |  |  |  |  |  |  |  |
| AIC | k<sub>1</sub> | p<sub>1</sub> |  |  |  |  | R<sup>2</sup> |
| 6537.8 | 0.4975 ± 0.1927 | **0.0099** |  |  |  |  | 0.0171 |
| **Interpret:** Pairwise comparisons showed significant differences between no and high, and between low and high uncertainty conditions on left hemisphere. |  |  |  |  |  |  |  |
| **Model 6:** betaPow ~ 1 + k<sub>1</sub>uncerID + 1\|subID, exclude stimID ≠ 1 \| laterID == 1 \| uncerID == 3 |  |  |  |  |  |  |  |
| AIC | k<sub>1</sub> | p<sub>1</sub> |  |  |  |  | R<sup>2</sup> |
| 6012.3 | 0.3342 ± 0.1950 | 0.0869 |  |  |  |  | 0.0306 |
| **Model 7:** betaPow ~ 1 + k<sub>1</sub>uncerID + 1\|subID, exclude stimID ≠ 1 \| laterID == 1 \| uncerID == 2 |  |  |  |  |  |  |  |
| AIC | k<sub>1</sub> | p<sub>1</sub> |  |  |  |  | R<sup>2</sup> |
| 5956.3 | 0.4149 ± 0.0953 | **1.4618 × 10<sup>-5</sup>** |  |  |  |  | 0.0328 |
| **Model 8:** betaPow ~ 1 + k<sub>1</sub>uncerID + 1\|subID, exclude stimID ≠ 1 \| laterID == 1 \| uncerID == 1 |  |  |  |  |  |  |  |
| AIC | k<sub>1</sub> | p<sub>1</sub> |  |  |  |  | R<sup>2</sup> |
| 6419.1 | 0.5120 ± 0.1851 | **0.0058** |  |  |  |  | 0.0054 |
| **Interpret:** Pairwise comparisons showed significant differences between no and high, and between low and high uncertainty conditions on right hemisphere. |  |  |  |  |  |  |  |
| **Model 9:** uncerID ~ 1 + k<sub>1</sub>betaPow + k<sub>2</sub>alphaPow + 1\|subID, exclude stimID ≠ 1 \| laterID == 2, only consider power modulation on the left hemisphere. |  |  |  |  |  |  |  |
| AIC | k<sub>1</sub> | p<sub>1</sub> | k<sub>2</sub> | p<sub>2</sub> |  |  | R<sup>2</sup> |
| 4233.4 | 0.0134 ± 0.0059 | **0.0239** | 0.0092 ± 0.0046 | **0.0471** |  |  | 0.0083 |
| **Model 10:** uncerID ~ 1 + k<sub>1</sub>betaPow + k<sub>2</sub>alphaPow + 1\|subID, exclude stimID ≠ 1 \| laterID == 1, only consider power modulation on the right hemisphere. |  |  |  |  |  |  |  |
| AIC | k<sub>1</sub> | p<sub>1</sub> | k<sub>2</sub> | p<sub>2</sub> |  |  | R<sup>2</sup> |
| 4222 | 0.0200 ± 0.0065 | **0.0021** | 0.0125 ± 0.0046 | **0.0062** |  |  | 0.0151 |
| **Compare (Model 9, Model 10)** |  |  |  |  |  |  |  |
| LRStat | deltaDF | pValue |  |  |  |  |  |
| 11.377 | 0 | **<0.0001** |  |  |  |  |  |
| **Interpret:** Modulation of beta and alpha power in both hemispheres can predict the level of uncertainty, with greater predictive accuracy observed when using modulation from the right hemisphere. |  |  |  |  |  |  |  |

betaPow=normalised beta power after the uncertainty cue; uncerID=uncertainty condition index, 1, 2, and 3 indicate no, low, and high uncertainty conditions, respectively; laterID=laterality condition index, 1 and 2 indicate the left and right hemispheres, respectively; inter=interaction; subID=subject index; stimID=rTMD condition index, 1, 2, and 3 indicate no, regular, and irregular rTMS conditions, respectively; AIC=Akaike information criterion; alphaPow=normalised alpha power after the uncertainty cue; LRStat= Likelihood Ratio Statistic; deltaDF=The difference in degrees of freedom between the models; pValue= statistical p-value indicating whether the improvement in fit is statistically significant or not. Each model included an independent random intercept to count for variability across participants. The inclusion of random slop(s) between the predictor(s) and the dependent variable was determined through model comparison. For each GLME model, the parameters were estimated based on maximum likelihood using Laplace approximation, the AIC, estimate value with standard error of the coefficient ( $k \pm SE$ ), pre-corrected $p$ -value ( $p$ ), and proportion
of variability in the response explained by the fitted model ( $R^2$ ) were reported. P-values survive multiple comparison correction following FDR approach were highlighted.

**Supplementary Table 3.** Generalized linear mixed-effects (GLME) modelling revealed no significant association between reaction time and the timing of rTMS pulses in the rTMS conditions.

|  |  |  |  |  |  |  |  |
| --- | --- | --- | --- | --- | --- | --- | --- |
| **Model 1:** $RT \sim 1 + k_1 \text{uncerID} + k_2 \text{lag} + k_{inter} \text{uncerID} * \text{lag} + 1 \vert \text{subID} + (-1 + \text{uncerID} \vert \text{subID})$, exclude $\text{stimID} == 1$ | | | | | | | |
| AIC | $k_1$ | $p_1$ | $k_2$ | $p_2$ | $k_{inter}$ | $p_{inter}$ | $R^2$ |
| -12507 | $0.1026 \pm 0.0201$ | **$3.5545 \times 10^{-7}$** | $0.0060 \pm 0.0458$ | 0.8959 | $-0.0510 \pm 0.0205$ | **0.0129** | 0.5430 |
| **Interpret:** Significant interaction between uncertainty and lag, therefore the model was updated by considering each individual uncertainty level. |  |  |  |  |  |  |  |
| **Model 2:** $RT \sim 1 + k_1 \text{lag} + 1 \vert \text{subID}$, exclude $\text{stimID} == 1 \mid \text{uncerID} \neq 1$ | | | | | | | |
| AIC | $k_1$ | $p_1$ | $R^2$ | | | | |
| -3498.8 | $0.0182 \pm 0.0434$ | 0.6756 | 0.1965 | | | | |
| **Model 3:** $RT \sim 1 + k_1 \text{lag} + 1 \vert \text{subID}$, exclude $\text{stimID} == 1 \mid \text{uncerID} \neq 2$ | | | | | | | |
| AIC | $k_1$ | $p_1$ | $R^2$ | | | | |
| -4387.2 | $-0.1256 \pm 0.0609$ | 0.0444 | 0.3216 | | | | |
| **Model 4:** $RT \sim 1 + k_1 \text{lag} + 1 \vert \text{subID}$, exclude $\text{stimID} == 1 \mid \text{uncerID} \neq 3$ | | | | | | | |
| AIC | $k_1$ | $p_1$ | $R^2$ | | | | |
| -4599 | $-0.0198 \pm 0.0531$ | 0.7092 | 0.3417 | | | | |
| **Interpret:** There is no significant association between lag and reaction time in rTMS conditions. |  |  |  |  |  |  |  |

RT=reaction time; uncerID=uncertainty condition index, 1, 2, and 3 indicate no, low, and high uncertainty conditions, respectively; lag=time lag/interval between the first TMS pulse and the go cue; inter=interaction; subID=subject index; stimID=rTMD condition index, 1, 2, and 3 indicate no, regular, and irregular rTMS conditions, respectively; AIC=Akaike information criterion. In each model, an independent random slope(s) between the predictor(s) and the dependent variable as well as an independent random intercept (s) were included. For each GLME model, the parameters were estimated based on maximum likelihood using Laplace approximation, the AIC, estimate value with standard error of the coefficient ( $k \pm SE$ ), pre-corrected $p$ -value ( $p$ ), and proportion of variability in the response explained by the fitted model ( $R^2$ ) were reported. P-values survive multiple comparison correction following FDR approach were highlighted.

**Supplementary Fig. 1.**
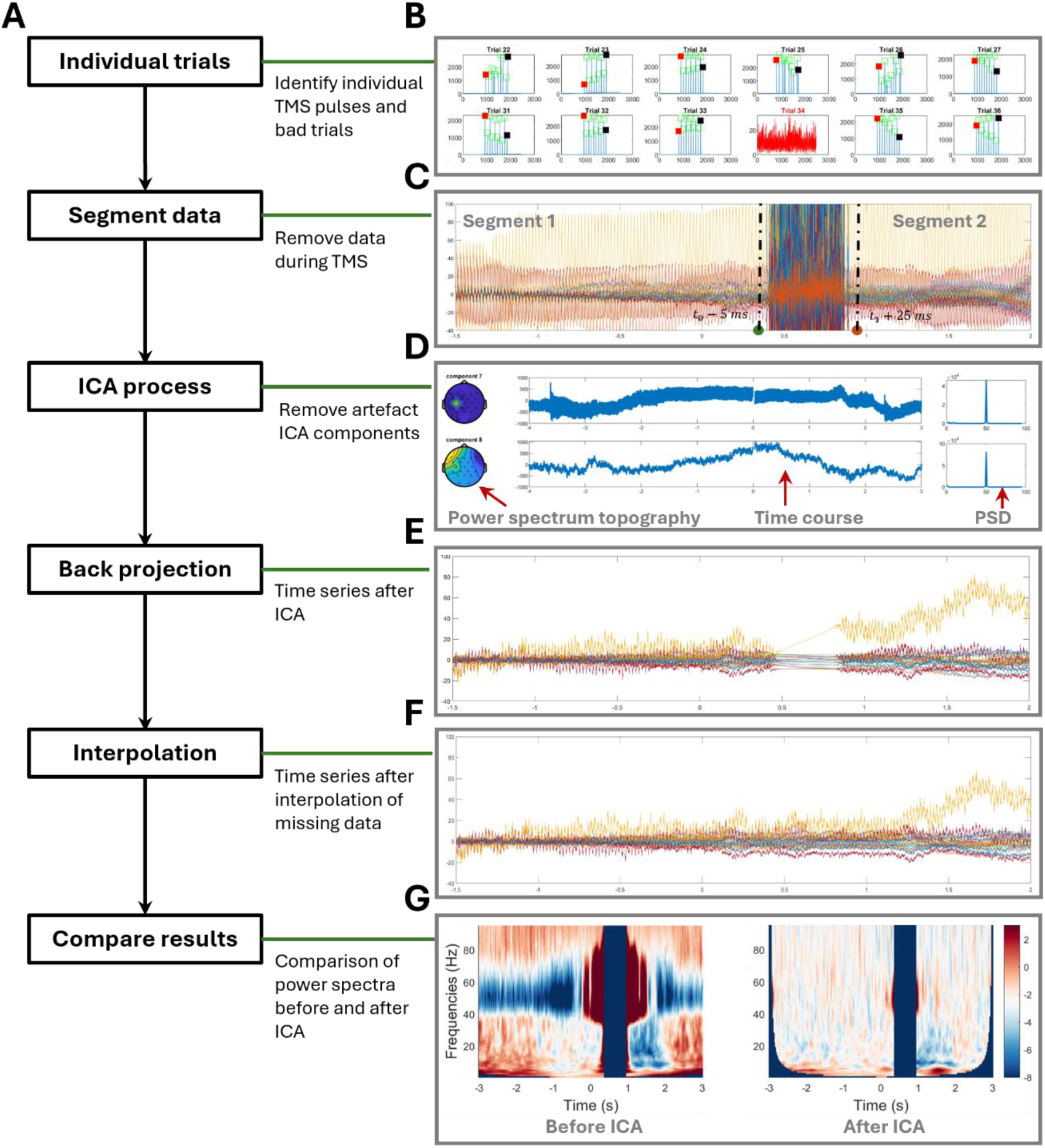
EEG data preprocessing pipeline. **(A)** Schematic of the preprocessing steps used in this study. **(B)** Identification of individual TMS pulses and exclusion of bad trials. **(C)** Removal of data from 5 ms before the first TMS pulse to 25 ms after the last TMS pulse. **(D)** Identification and removal of artefactual components using Independent Component Analysis. **(E)** Reconstruction of the time series from the cleaned components. **(F)** Interpolation of data segments missing during TMS pulses by mirroring the signal immediately before and after each pulse to ensure continuity. **(G)** Example comparison of power spectra before and after preprocessing.

**Supplementary Fig. 2.**
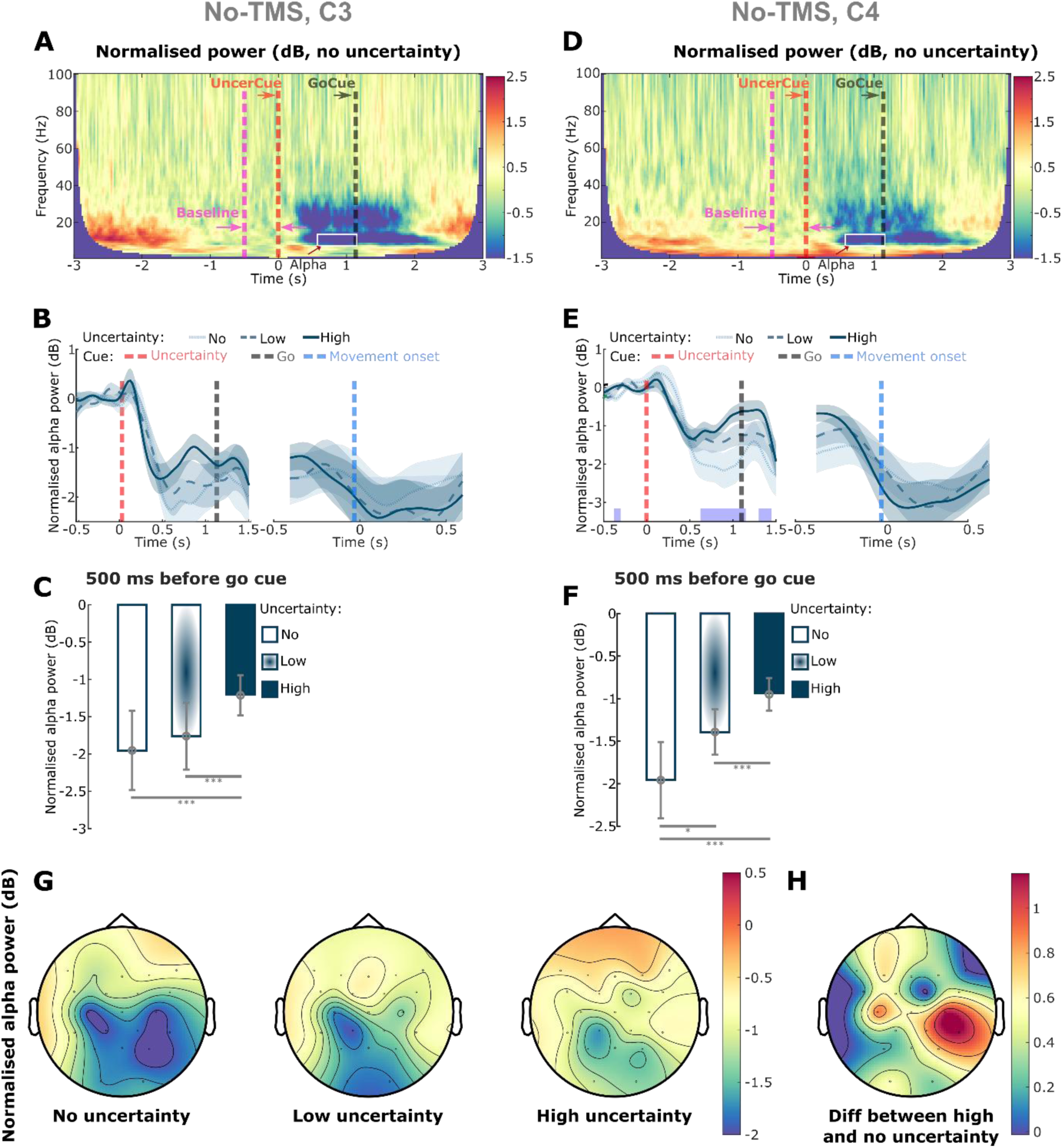
Bilateral modulation of alpha by uncertainty. **(A)** Group-averaged time-frequency power spectra from EEG channel C3 (left hemisphere), aligned to the onset of the ‘uncertainty’ cue (red dashed line), in the no-rTMS/no-uncertainty condition. Power spectra were normalised to a 500-millisecond pre-cue resting baseline for each trial. Alpha-band power (white box) showed modulations following the ‘uncertainty’ cue and prior to the ‘go’ cue (black dashed line). **(B)** Group-averaged time courses of alpha power, normalised to baseline, in different uncertainty conditions. Red, black, and blue dashed lines indicate the ‘uncertainty’, ‘go’, and ‘movement onset’ cues, respectively. **(C)** Comparison of alpha power modulation across different uncertainty conditions. Power was quantified as the average within 500-millisecond window preceding the ‘go’ cue, normalised (in dB) to the 500-millisecond pre-‘uncertainty’ cue baseline. Error bars represent the mean ± SEM across participants. **(D)-(F)** Same analyses as in (A)-(C), but for EEG channel C4 (right hemisphere). **(G)** Topographical map of alpha power modulation under conditions of no (left), low (middle), and high (right) uncertainty. Power was quantified as in (C) and (F). **(H)** Difference in alpha power modulation between high and no uncertainty conditions. *P < 0.05, **P < 0.01, ***P < 0.001. P-values in (C) and (F) were derived from generalised linear mixed-effects models applied to individual trials and corrected for multiple comparisons using FDR. Purple bars in (B) and (E) indicate significant differences between no- and high-uncertainty conditions (cluster-based permutation test).

**Supplementary Fig. 3.**
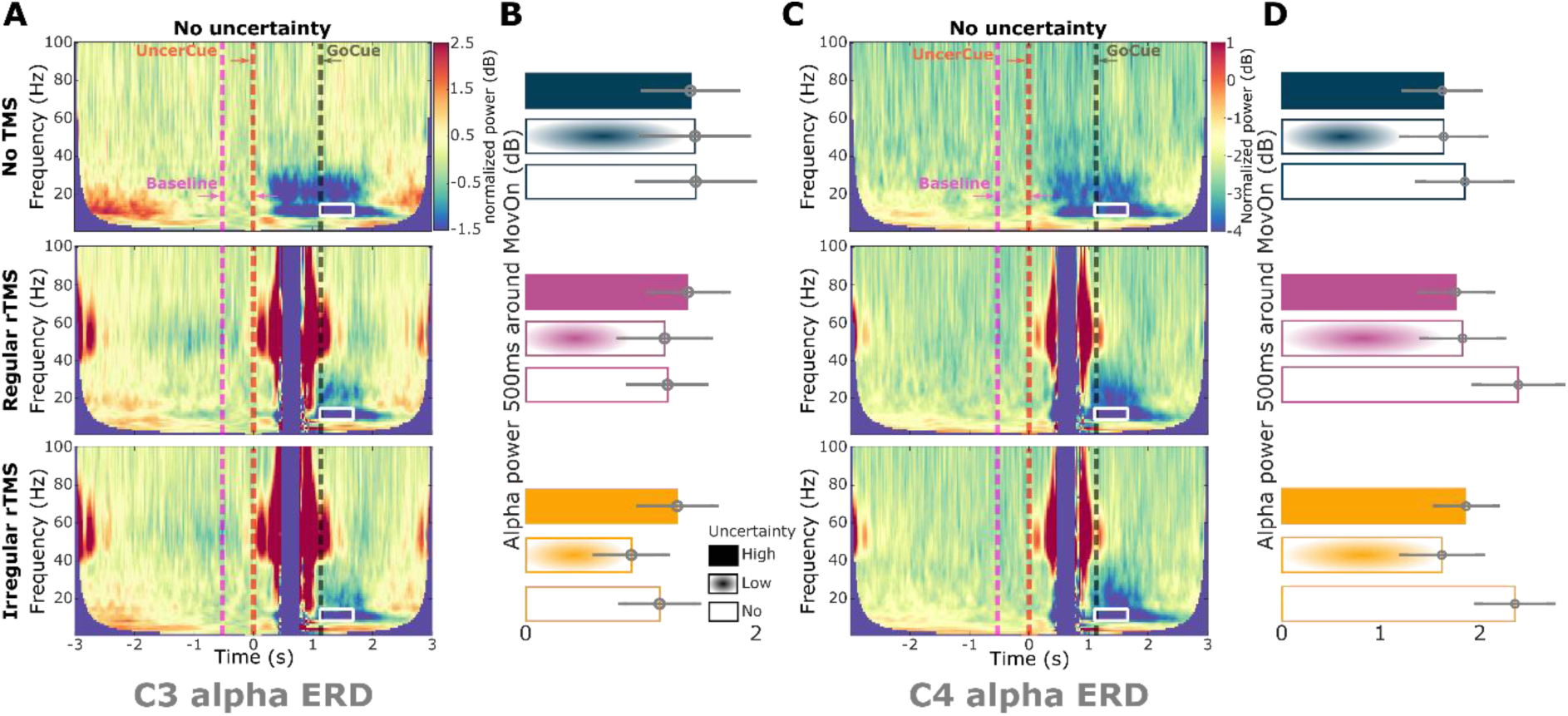
Alpha ERD was not modulated with rTMS. **(A)** Group-averaged time-frequency power spectra from EEG channel C3 (left hemisphere), aligned to the onset of the ‘uncertainty’ cue (red dashed line), shown for three conditions: no uncertainty and no TMS (up), regular rTMS (middle), and irregular rTMS (bottom). Power spectra were normalised to a 500-millisecond pre-cue resting baseline (pink dashed line) for each trial. Alpha-band activity (white box) showed modulation following movement onset (black dashed line). **(B)** Comparison of alpha ERD across different uncertainty and rTMS conditions. Power was quantified as the average within a 500-ms window preceding the ‘uncertainty’ cue and normalised (in dB) to the 500 ms window centered at movement onset. Error bars represent mean ± SEM across participants. **(C)-(D)** Same analyses as in (A)-(B), but for EEG channel C4 (right hemisphere). *P < 0.05. P-values in (B) and (D) were obtained using generalised linear mixed-effects models applied to individual trials and corrected for multiple comparisons using FDR.

**Supplementary Fig. 4.**
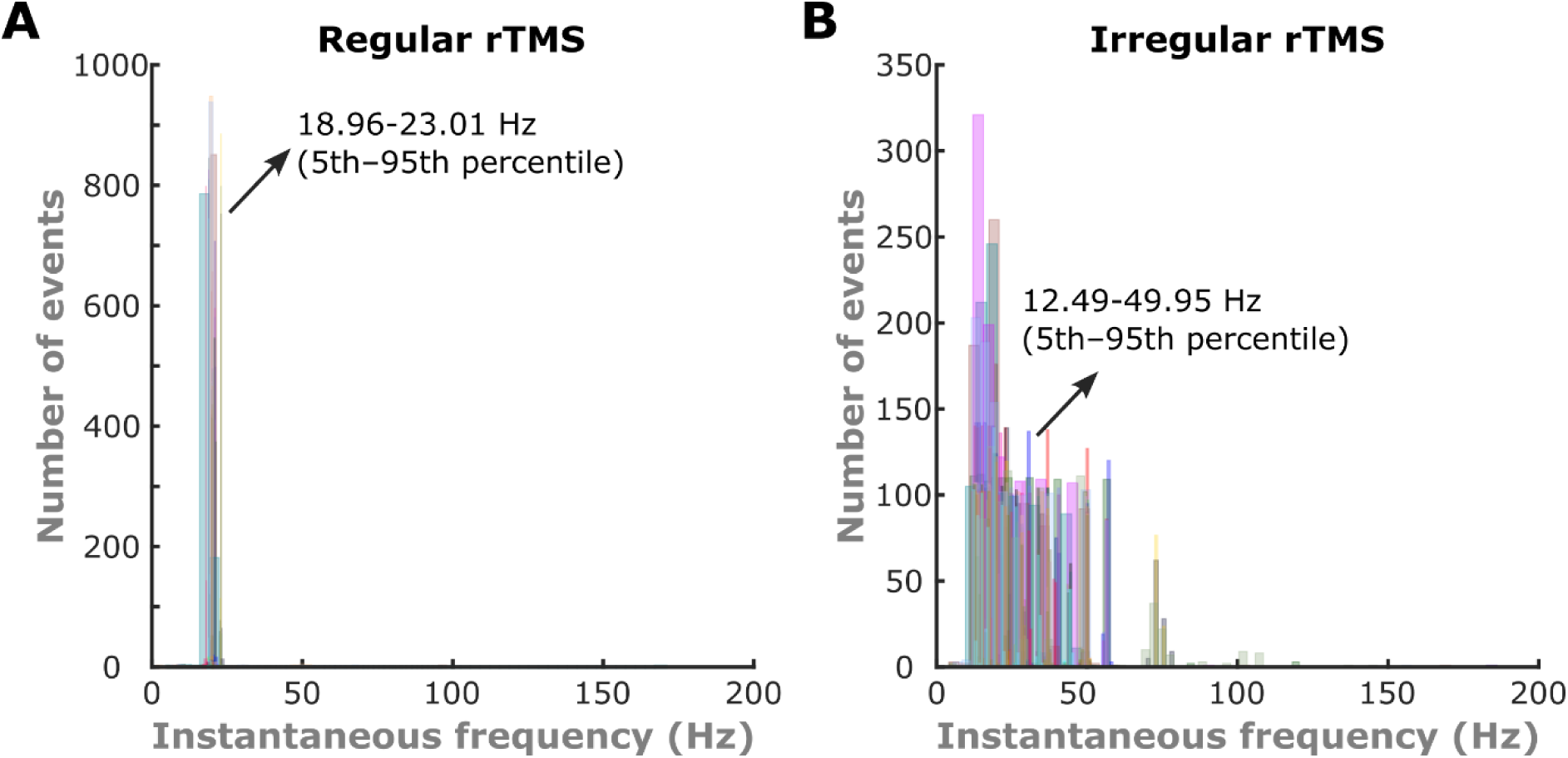
Histogram of instantaneous TMS pulse frequencies in the regular and irregular rTMS conditions. **(A)** In the regular rTMS condition, instantaneous frequencies ranged from 18.96 to 23.01 Hz (5th–95th percentile). **(B)** In the irregular rTMS condition, instantaneous frequencies ranged from 12.49 to 49.95 Hz (5th–95th percentile). Instantaneous frequency was computed by identifying the timing of each TMS pulse, calculating inter-pulse intervals, and converting these intervals to frequencies.

